# Climate, population size, and dispersal influences mutational load across the landscape in *Vitis arizonica*

**DOI:** 10.1101/2024.08.22.609253

**Authors:** Christopher J. Fiscus, Jonás A. Aguirre-Liguori, Garren R. J. Gaut, Brandon S. Gaut

## Abstract

The interplay between genetics and the environment determines a population’s ability to survive. Plants, being immobile, are particularly vulnerable to environmental shifts and must adapt to their local environment or face extinction. While considerable efforts have been devoted to identifying adaptive genetic variation and to model its relationship with climate, the role of deleterious variation has been largely overlooked. To address this gap, we studied the landscape genomics of *Vitis arizonica*, a grape species endemic to the American Southwest and a crop wild relative of the domesticated grapevine. We estimated mutational load, a component of genetic load, in 162 individuals sampled across the present species range and built temporal species distribution models (SDMs) to project the historical, present, and future distributions of *V. arizonica* to infer species’ range dynamics. Mutational load was highest for individuals at an inferred leading edge of range expansion. Using random forest regression (RF) models, we examined the relationship between mutational load and climatic variation. The RF models, which we transformed by a weighting method to account for correlated predictors, identified climatic variables, historical dispersion distances, and heterozygosity as high-ranking features. Our findings show that mutational load can be predicted and identifies features that contribute to load. These results provide a foundation for integrating mutational load into broader efforts to understand species adaptation and maladaptation in the face of climate change.

**SIGNIFICANCE STATEMENT:** The accumulation of deleterious genetic variation (i.e., genetic load) in a genome can reduce organismal fitness and hinder adaptation to local conditions. While the genetic mechanisms contributing to genetic load have been well-studied, its interaction with environmental variation is less understood. Here we explored the relationship between climatic variation, species range, and genetic load in *Vitis arizonica*, a wild grape species native to the American Southwest. We identified associations between climatic variation and genetic load and built a machine learning model to predict genetic load under future climate scenarios. Our findings suggest that the species range will expand and that genetic load will slightly increase at the population level by the end of the century. This work enhances our understanding of the environmental factors influencing genetic load.

## INTRODUCTION

Mutations are the raw material that fuels evolutionary change, and they represent a range of effects on fitness. A small proportion of new mutations confer a fitness advantage and many are evolutionarily neutral. However, the vast majority are deleterious (Eyre-Walker & Keightley 2007), contribute to genetic load despite often being only transient in the population due to selection against them, and thus affect evolutionary outcomes (Hedrick & Kalinowski 2000; Robinson et al. 2023). Surveying the number, prevalence, and fitness effects of deleterious mutations is crucial for understanding a wide array of biological phenomena, ranging from the incidence and severity of genetic diseases (Kryukov et al. 2007) to the conservation status of wild populations (Kyriazis et al. 2021).

Given their importance and ubiquity, deleterious variants have been the focus of many theoretical and empirical studies (Robinson et al. 2023). These studies have shown that the frequency and number of deleterious variants within a population are shaped by several evolutionary parameters, including effective population size (*Ne*) (Charlesworth & Charlesworth 1998). Populations with lower *Ne* tend to accumulate more deleterious mutations than populations with higher *Ne*, due to increased genetic drift and reduced selection efficacy.

Accordingly, the number and frequency of deleterious variants often increase through population bottlenecks and other demographic events (Lohmueller 2014). However, the accumulation of deleterious variants also depends on factors beyond *Ne*, such as dominance coefficients (Simons et al. 2014), the distribution of fitness effects (DFE), and the duration and timing of a bottleneck (Brandvain & Wright 2016; Bortoluzzi et al. 2020). In fact, bottlenecks of sufficient duration and severity can purge highly deleterious, homozygous mutations (Grossen et al. 2020; Femerling et al. 2023). Nonetheless, demography helps explain the increased number and frequencies of deleterious variants in domesticated species (Renaut & Rieseberg 2015; Liu et al. 2017; Moyers et al. 2018) and in small populations of conservation concern (Femerling et al. 2023). The accumulation of deleterious variants can also be affected by hitchhiking (i.e., linkage load) and gene flow. Gene flow can either increase or decrease genetic load depending on the complement of deleterious variants on introgressed haplotypes (Kim et al. 2018; Zhang et al. 2020; Xiao et al. 2023).

The evolutionary processes that shape genetic load do not act uniformly across geographic landscapes. Populations at range edges often experience environmental marginality, resulting in smaller *Ne* at the edges of ecological or geographic niches (Angert et al. 2020) and the accumulation of deleterious variants in edge populations. This expectation may differ, however, between ‘leading’ and ‘trailing’ edges of a range shift (Angert et al. 2020). Populations at the leading (i.e. expanding) edges of a range shift are more likely to represent serial founder events, assortative mating and strong selection pressure to adapt to new biotic and abiotic environments. Moreover, once deleterious mutations are established, they can propagate with an expanding range front via gene surfing (Travis et al. 2007; Excoffier et al. 2009). Consistent with these expectations, increased numbers and frequencies of deleterious variants have been found at range edges in both laboratory (Bosshard et al. 2017; Weiss-Lehman et al. 2017) and natural populations (González-Martínez et al. 2017; Willi et al. 2018; Rougemont et al. 2020; Takou et al. 2021; Willi et al. 2022).

Because populations at the leading-edge of range expansion often encounter new or unique environmental challenges, it seems reasonable to posit that the number and frequency of deleterious variants co-varies with environmental and climatic markers. Although leading-edge effects have been implicated in reduced rates of adaptation (Sánchez-Castro et al. 2022) and shifts in mating system, there have been few attempts to relate distributions of deleterious variants to climatic variation (Willi et al. 2022). We believe that exploring the connection between deleterious variants and climate is worthwhile for at least two reasons.

First, such an exploration is likely to provide additional insights into the evolutionary processes that shape the spatial distribution of genetic variation. Second, since deleterious variants affect fitness, understanding their patterns can inform about the probability of population persistence (Aguirre-Liguori et al. 2021) and adaptation in the context of climate change (Sánchez-Castro et al. 2022). Recently, substantial attention has focused on modeling the fate of putatively adaptive mutations to predict the fate of populations under predicted climate change (Fitzpatrick & Keller 2015; Waldvogel et al. 2020; Capblancq, Fitzpatrick, et al. 2020). Except in the special case of antagonistic pleiotropy, these approaches typically overlook deleterious variants, representing a potentially major conceptual gap in the field of climate genomics (Aguirre-Liguori et al. 2021).

Here we examine deleterious variation and evaluate its relationship to features such as *Ne*, dispersal, and climate across a landscape. To achieve this, we measured deleterious variants in individuals across the geographic range of a wild plant species, *Vitis arizonica*, which is a perennial crop wild relative (CWR) of the domesticated grapevine (*V. vinifera* ssp *vinifera*). CWRs have multi-billion dollar impacts on the global economy (Bohra et al. 2022), because they provide agronomically beneficial traits as rootstocks or through hybrid breeding. CWRs are of urgent conservation concern (Khoury et al. 2020), particularly as climate change initiates species’ range shifts. *V. arizonica* is an interesting candidate for conservation efforts for two reasons. First, it is native to a wide environmental range encompassing Northern Mexico and the U.S. Southwest (Heinitz et al. 2019), where extreme heat and drought are common.

Second, it has the potential to contribute valuable agronomic traits – such as drought tolerance, salinity tolerance, and pathogen resistance – to viticulture. In fact, *V. arizonica* segregates for resistance to *Xylella fastiodisa* (Riaz et al. 2020, 2018; Morales-Cruz et al. 2023), a plant pathogen that causes Pierce’s Disease (PD) in grapevines and that also infects major crops like almonds, olives, and coffee. Consequently, *V. arizonica* has already been utilized in breeding programs to introgress PD-resistance into *V. vinifera* varieties (Quinton 2019).

In this study, we estimate mutational load, a measure of genetic load, for a sample of ∼162 resequenced *V. arizonica* individuals. We then compile a collection of genomic, geographic, and bioclimatic features corresponding to each sampling location and examine associations between these features and mutational load. Using this array of data, we address three key questions: First, what is the distribution of putatively deleterious variants within *V. arizonica* individuals, and how does this distribution vary across the sampling range? Second, which aspects of population history and environmental conditions best predict the distribution of deleterious variants across the landscape? Third, can a predictive model be used to project future changes in mutational load under different climate scenarios?

## RESULTS

### Mutational load estimates across the landscape

We called variants in a cohort of 172 resequenced individuals sampled across much of the native range of *V. arizonica* (Morales-Cruz et al. 2023) (**Table S1**). After site and sample filtering (Materials and Methods, **Figure S1**), the final dataset represented genotypes for 162 individuals across 1,320,747 biallelic SNPs with a minor allele frequency of 1% or greater. The filtered dataset had low levels of missing data, averaging 0.39% missing calls both per site and per sample. We annotated these SNPs for predicted effects on protein function, identifying 140,801 synonymous (sSNPs) and 146,830 nonsynonymous SNPs (nSNPs). The latter included 40,145 putatively deleterious SNPs (dSNPs), as predicted by SIFT-4G (Vaser et al. 2016). We polarized sSNPs and nSNPs using six individuals from two congeneric species (*V. girdiana* and *V. monticola*). After applying our polarizing criteria (Materials and Methods), we assigned ancestral and derived alleles for 53.02% (74,656 / 140,801) of sSNPS, for 54.01% (79,301 / 146,830) of nSNPs, and for 55.89% (22,436 / 40,145) of dSNPS.

We expected both nSNPs and dSNPs to have been subject to purifying selection and thus segregate at lower frequencies compared to sSNPs. We tested this hypothesis by calculating the site frequency spectrum (SFS) for each SNP class across the entire sample **(Figure 1A)**. As expected, the SFS for nSNPs was significantly different than that of sSNPs (Kolmogorov-Smirnov test, D = 0.047, p < 2.2e-16), reflecting an enrichment for low frequency derived nSNP alleles (Chi-squared test, χ^2^ = 367.96, df = 1, p < 2.2 X 10^-16^). The SFS for dSNPs was also significantly different than the SFS for nSNPs (Kolmogorov-Smirnov test, D = 0.033, p = 6.66 X 10^-16^), with an even greater enrichment of low frequency variation compared to nSNPs (Chi-squared test, χ^2^ = 96.03, df = 1, p < 2.2 X 10^-16^). Thus, nSNPs and dSNPs segregated as expected, with the SFS dominated by low frequency variants.

**Figure 1.**
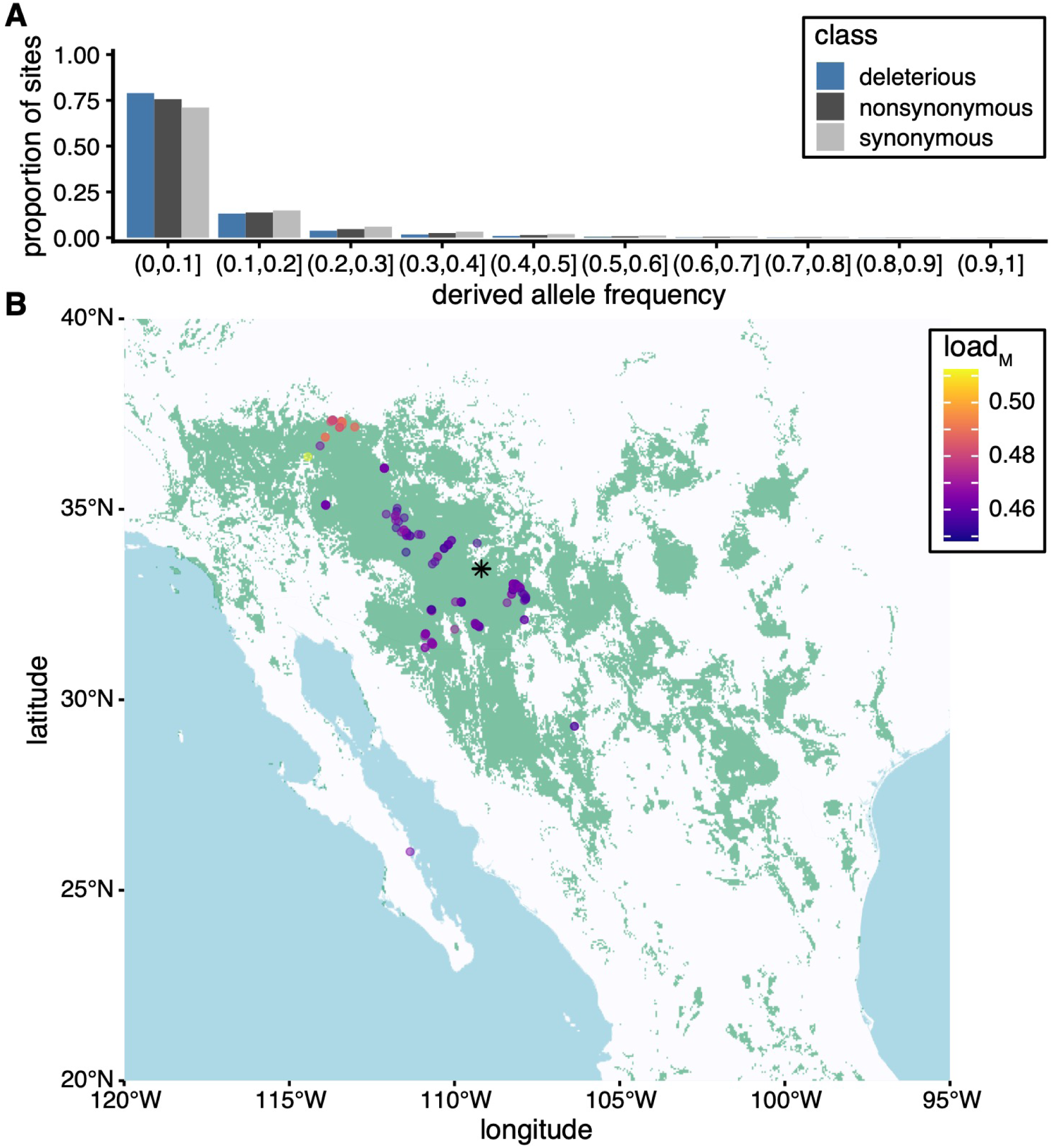
Site frequency spectra and mutational load in *V. arizonica*. (A) Derived allele frequency for three SNP classes. (B) Present species distribution model (green) based on WorldClim (bioclimatic averages from 1970 to 2000) and GBIF species occurrence data. The points represent sampling locations for individuals used in genetic analyses and are colored according to the load_M_ estimate per individual. The black asterisk indicates the geographic centroid of the predicted range.

We next estimated mutational load per individual (load_M_) following a previous method (Willi et al. 2018). Briefly, load_M_ measured the proportion of derived alleles at polarized nSNP sites divided by the sum of two values: the proportion of derived alleles at polarized nSNP sites and the proportion of derived alleles at polarized sSNP sites (Materials and Methods). The estimates of load_M_ based on nSNPs varied 11% between individuals and were distributed unevenly across sampling locales, with higher values common in the northern extreme of the predicted species range (**Figure 1B**). To complement our estimate of load_M_ based on nSNPs, we also calculated load_M_ based on the frequency of dSNPs. The two load_M_ measures were highly correlated (R^2^ = 0.73, p < 2.2 X 10^-16^, **Figure S2**) and yielded qualitatively identical results for downstream analyses. Thus, for simplicity, we opted to report the measure based on nSNPs, as with previous work (Willi et al. 2018).

### Compilation of features to predict load_M_

A central goal was to build models to evaluate the relative contributions of genetics, the environment, and species’ range dynamics for predicting load_M_. With this objective in mind, we compiled a set of 24 features related to genomic diversity, climatic diversity, and species’ range for each individual (**Table 1**, **Figure S3**). The features included the 19 WorldClim 2 bioclimatic variables, which summarize historical climate variation from 1970 - 2000 (Fick & Hijmans 2017) at each collection site. To represent genomic variation, we calculated observed heterozygosity (H_O_) for each individual, reasoning that this measure reflects aspects of historical *Ne* (Crow & Kimura 1970).

**Table 1.** Summary of Predictors.

| Feature | Class | Description |
| --- | --- | --- |
| Ho | genetic | Observed heterozygosity |
| bio1-bio19 | bioclimatic | WorldClim bioclimatic variables |
| distance from geographic centroid | geography | distance from geographic centroid based on species distribution modeling |
| distance from range edge | geography | distance to nearest species distribution edge |
| distance from niche centroid | bioclimatic | distance from centroid of bioclimatic PCA |
| dispersal distance | dispersal | retroactive predicted dispersal distance from Last Glacial Maximum to the present |

The remaining four features were related to the species’ range, such as the distance of each individual to the nearest range edge or to the geographic or niche centroids. To calculate features based on species range dynamics, we built species distribution models (SDMs) that predicted the species’ range based on species occurrence (Occdownload Gbif.Org 2020). The SDMs were based on bioclimatic data from the present era (i.e., 1970 - 2000) and for the Last Glacial Maximum (LGM), ∼22,000 years ago **(Figure 2A)**. The present-day SDM was used to calculate the minimum geographic distance of each individual from the range edge, from the geographic centroid, and from the niche centroid (Materials and Methods).

**Figure 2.**
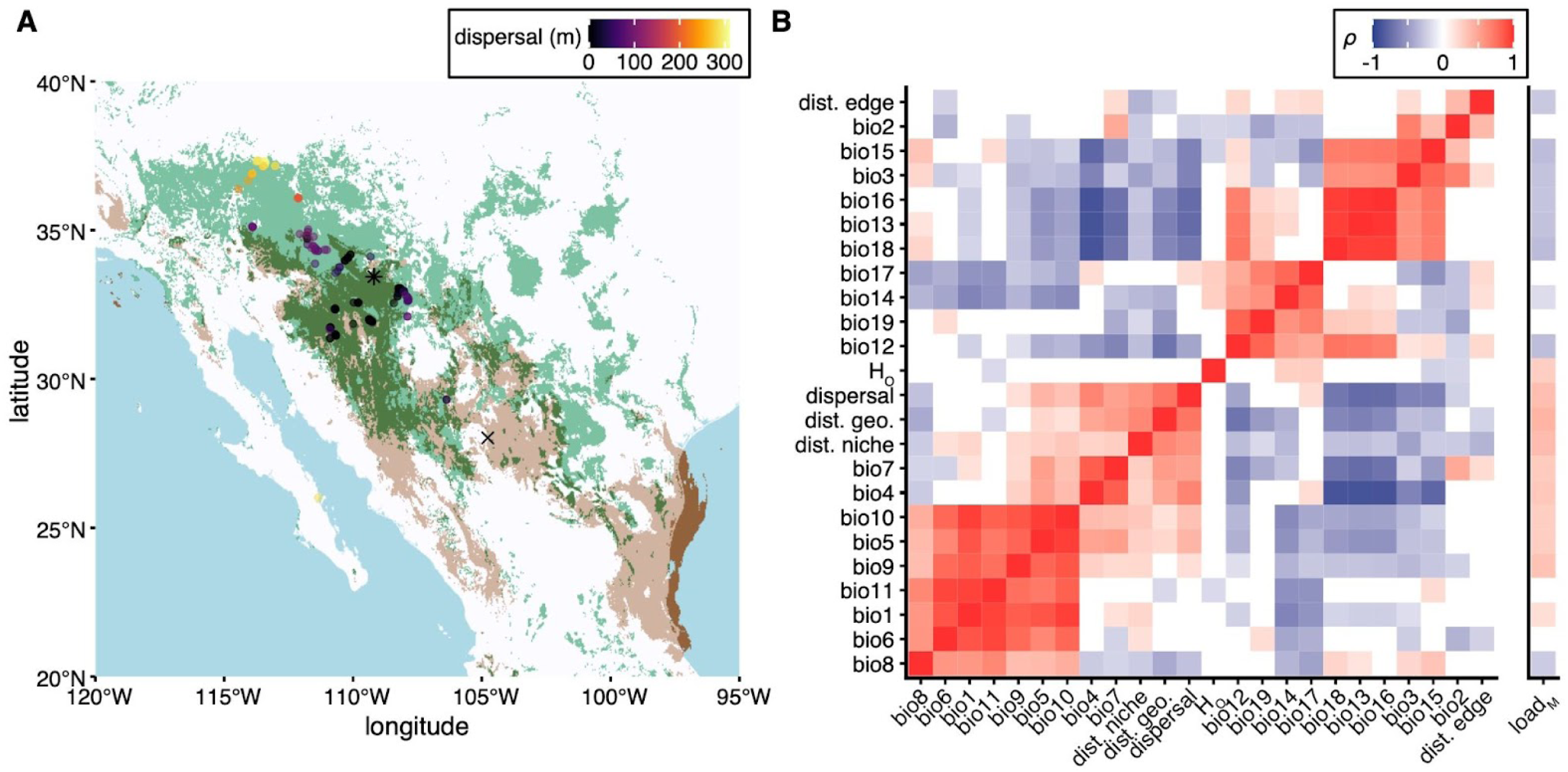
Sample dispersal and feature correlations. (A) Projected species distribution models (SDM) for the Last Glacial Maximum (∼22 KYA, brown), for the present (green), and for the overlap between SDMs (dark green). LGM SDM overlapping the Gulf of Mexico represents potential range during the LGM when sea levels were lower. Each point represents the sampling location for an individual and is colored according to predicted dispersal from the LGM to the present. The asterisk indicates the geographic centroid in the present, while the X indicates geographic centroid of the past. (B) Spearman’s correlations between features used in the RF model and between load_M_ and all features. In addition to the 19 WorldClim bioclimatic variables, the features include distance from the present-day SDM edge (dist. edge), observed heterozygosity (H_0_), the estimated dispersal distance shown in panel A (dispersal), the distance from the present-day geographic centroid (dist. geo), and the distance from the present-day niche centroid (dist. niche). Only correlation tests with p < 0.05 are filled.

A comparison of the two SDMs suggested that the species distribution moved and expanded over geologic time, with a shift in the geographic centroid by 736 km (34 degrees west of north) and a 40.71% expansion by area from the LGM to the present. However, the two SDMs overlapped for only 28.08% of the estimated present range, suggesting that there may have been substantial historical migration and dispersal. To estimate dispersal for our sample, we calculated the minimum distance from the present-day sampling location to the LGM SDM range edge. This approach was straightforward for individuals collected in locations constituting the present but not past range. For individuals collected at locations that were present in both SDMs, we assumed a dispersal distance of zero, despite climatic differences between the two epochs. Using these criteria, 84 sampling locations represented potential historical dispersions since the LGM, with a median distance of 89.97 km; another 78 individuals were located in overlapping regions that had no need for historical dispersal based on the inferred SDMs. The estimated dispersal distance was particularly pronounced for individuals sampled in more northern latitudes (**Figure 2A**), suggesting that this region represents a leading edge of range expansion.

The collection of genetic, geographic, and climatic features constitute a multivariate dataset that can be used to predict load_M_. As is common with such datasets, however, the features exhibited a complex pattern of correlated relationships (**Figure 2B)**. For example, bioclimatic variables related to temperature (e.g. bio1, bio5, bio6, bio8, bio9, bio10, and bio11) were positively correlated with each other, as were precipitation variables (e.g. bio13, bio17 and bio18). Other bioclimatic variables had negative correlations, including temperature variability (bio4 and bio7) relative to precipitation measures. Geographic measures also exhibited complex relationships, e.g., the dispersal distance and the distances to geographic and niche centroids were significantly correlated with one another but negatively correlated with most precipitation variables. In short, few of the features were statistically independent and thus evaluation of potential predictors of load_M_ must account for complex correlative relationships.

### Using random forest regression models to predict load_M_ in the present

We built random forest (RF) regression models to predict load_M_ using genetic, bioclimatic, and geographic features as predictors. We chose RF for its ability to infer non-linear relationships between predictors and response variables, which have been commonly observed when modeling relationships between climatic and genetic variation (Aguirre-Liguori et al. 2017; Capblancq & Forester 2021), and because it provides a straightforward measure of the importance of each feature to the model’s predictive performance. However, feature importance can be affected by correlations between predictors. For instance, when two features are perfectly correlated the importance attributed to either feature is assigned randomly to one, making it appear that the second is not relevant for predicting the response (Breiman 2001). In the RF framework, feature importance is a valid measure of a feature’s contribution to the model (i.e. the inclusion of the second, perfectly-correlated feature does not provide the model with any new information), but these importances may not accurately reflect the biological relevance of the predictor to the response.

Since our goal was to be able to interpret the biological relevance of features for predicting load_M_, we applied Johnson’s relative weights (Johnson 2000) to RF regression models. Johnson’s method consists of first producing an orthogonal approximation of the data by calculating the singular value decomposition (SVD), then using the SVD component vectors as the predictors for the model. After training, the transformation parameter λ, which is calculated from the SVD, is used to project the orthogonal components’ feature importance back to the original feature space. We refer to this projected feature importance as the ’relative weighted feature importance’. With this transformation, one can calculate the relative weighted importance of each predictor, and the model output can be interpreted in the context of the original predictor features.

We trained RF models to predict load_M_ using both the original data and the orthogonally approximated data (using Johnson’s method) and compared the outputs of the two models. The RF model with the original data had an ‘out of bag’ R^2^ estimate of 0.68 (+/- 0.08 SE, as determined by 10-fold cross validation) and predicted the withheld test data with an R^2^ = 0.71 (**Figure 3A**, **Figure S4**). Of the 24 features, 10 jointly encompassed 90% of the cumulative relative importance (**Figure 3B**). The two most important features were dispersal distance and bio18 (precipitation of the warmest quarter); other top features included distance from geographic centroid, H_O_ and additional bioclimatic variables (bio3, bio4, bio8, bio13, bio15, and bio16). The explanatory power was slightly lower for the model built using transformed data; the ‘out of bag’ R^2^ estimate was 0.59 and the model yielded an R^2^ of 0.61 on the withheld test dataset, falling just below the interval of R^2^ = 0.70 +/- 0.08 (SE), as determined by 10-fold cross-validation (**Figure 3C**, **Figure S5**). After transformation, 90% of the cumulative relative weighted importance was explained by 15 predictors (**Figure 3D**). Eight of these predictors were also included in the set of 10 features in the original model but others (bio17, bio14, distance to niche centroid, distance to geographic edge, bio7, and bio12) were not important in the untransformed model. Notably, >50% of the relative cumulative importance in the weighted model was explained by the combination of bio8 (mean temperature of wettest quarter) and distance from geographic centroid. The estimated dispersal distance and observed heterozygosity were also important predators of load_M_.

**Figure 3.**
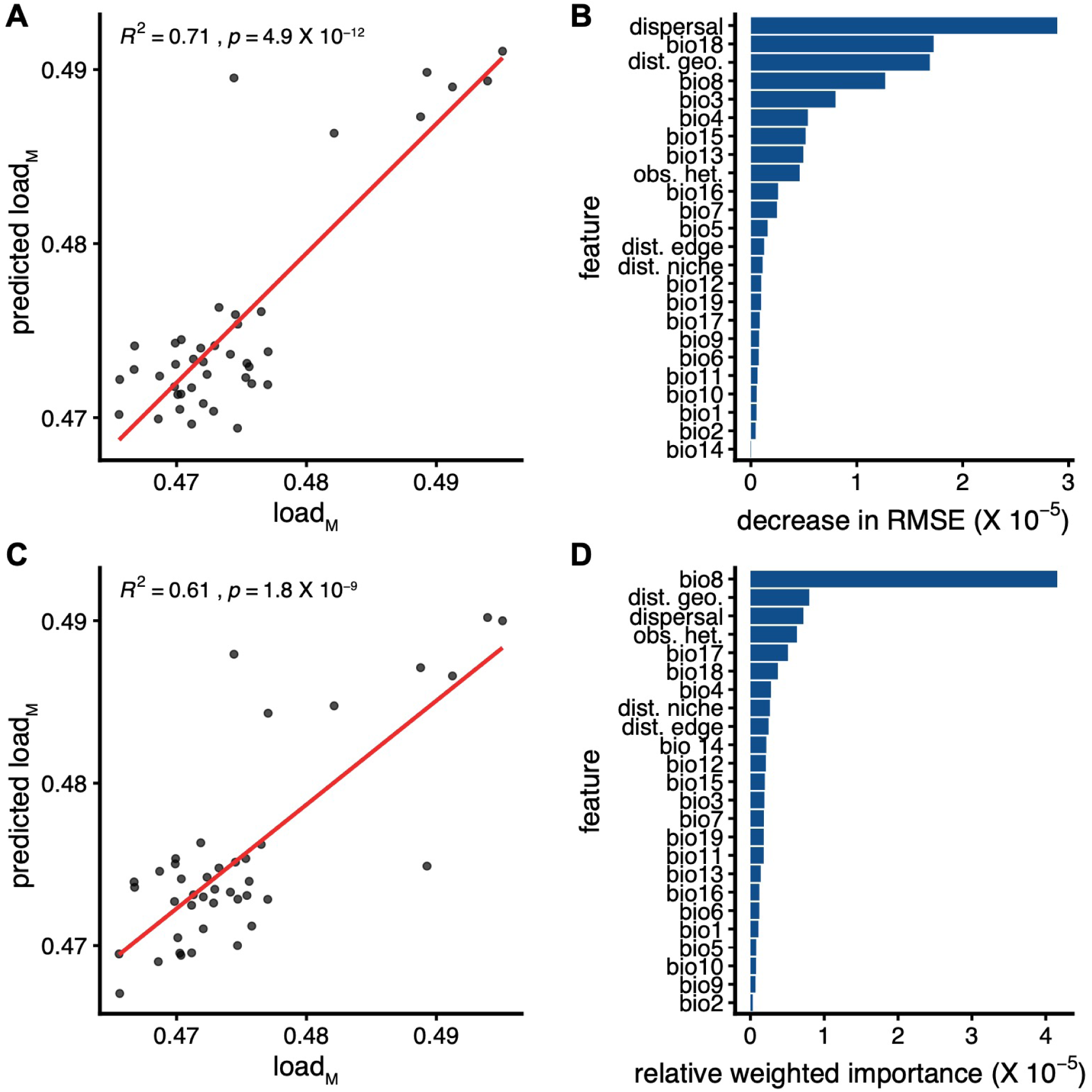
Random forest regression models to predict load_M_. Performance (A & C) and feature importance (B & D) of RF to predict load_M_. A and B correspond to the model fitted with untransformed data, while C and D correspond to the model fitted with data transformed by the Johnson method. In A and C, red lines indicate linear model fit. In B and D, the features are ranked by their inferred importance.

One concern is that the RF models do not explicitly account for the non-independence of observations due to relatedness between individuals (i.e., population structure). We addressed this concern in two ways. First, we fit univariate linear mixed models that included a relatedness (kinship) matrix to account for population structure. In total, 54.17% (13 / 24) of features were significantly associated with load_M_ in the linear mixed models (Bonferroni-adjusted p < 0.05, Table S2). The 13 associated features included all 10 (76.9%) of the most important features from the RF model using the natural data and 78.57% (11 / 14) of the most important features from the RF model using transformed data. Thus, the different modeling frameworks were complementary with respect to identifying explanatory features, even when relatedness was included. Second, we reasoned that the climatic data likely captured aspects of genetic relatedness, as individuals sampled in close proximity were likely to inhabit similar climates and share many alleles identical by descent. We tested this hypothesis by calculating the correlation between genetic distance and climatic distance matrices. The two matrices were significantly correlated (Mantel’s r = 0.41, 1,000 replicates, p = < 0.001), suggesting that the climatic features used in the RF models at least partially capture genetic relatedness.

### Species range and load_M_ at the end of the century

load_M_ can be reasonably well predicted using the compiled features that reflect present-era climate. But, what will happen to genetic load over time? Given that rapid anthropogenic-driven climate change is reshaping patterns of genotypic (Exposito-Alonso et al. 2022) and phenotypic (Saladin et al. 2020) diversity, we examined predicted climate trajectories, estimated a new dataset of features based on projected climates, and then applied our trained RF model to forecast load_M_ into the future. In total, we built SDMs for 16 potential climate trajectories, combining four Earth System Models (ESM) and four Shared Socioeconomic Pathway (SSP) scenarios, using forecasted climate averages from 2080 to 2100. The SSPs correspond to different levels of global greenhouse gas emissions and are defined by the cooperation between nations to curb these emissions. We considered four SSPs: SSP126, SSP245, SSP370, and SSP585, corresponding to CO_2_ emissions reaching net zero by 2050, net zero by 2075, remaining unchanged until 2050 then near net zero by 2100 or doubling by 2100, respectively (Riahi et al. 2017).

Overall, the future SDMs predicted that the potential species range is likely to shift northward and expand in size by 2100. Even under the most optimistic emissions scenario (i.e., SSP126, sustainability), the geographic centroid is predicted to move 87.80 to 226.45 km from the present centroid, with the degree of range shift dependent on the ESM (**Figure 4A**).

**Figure 4.**
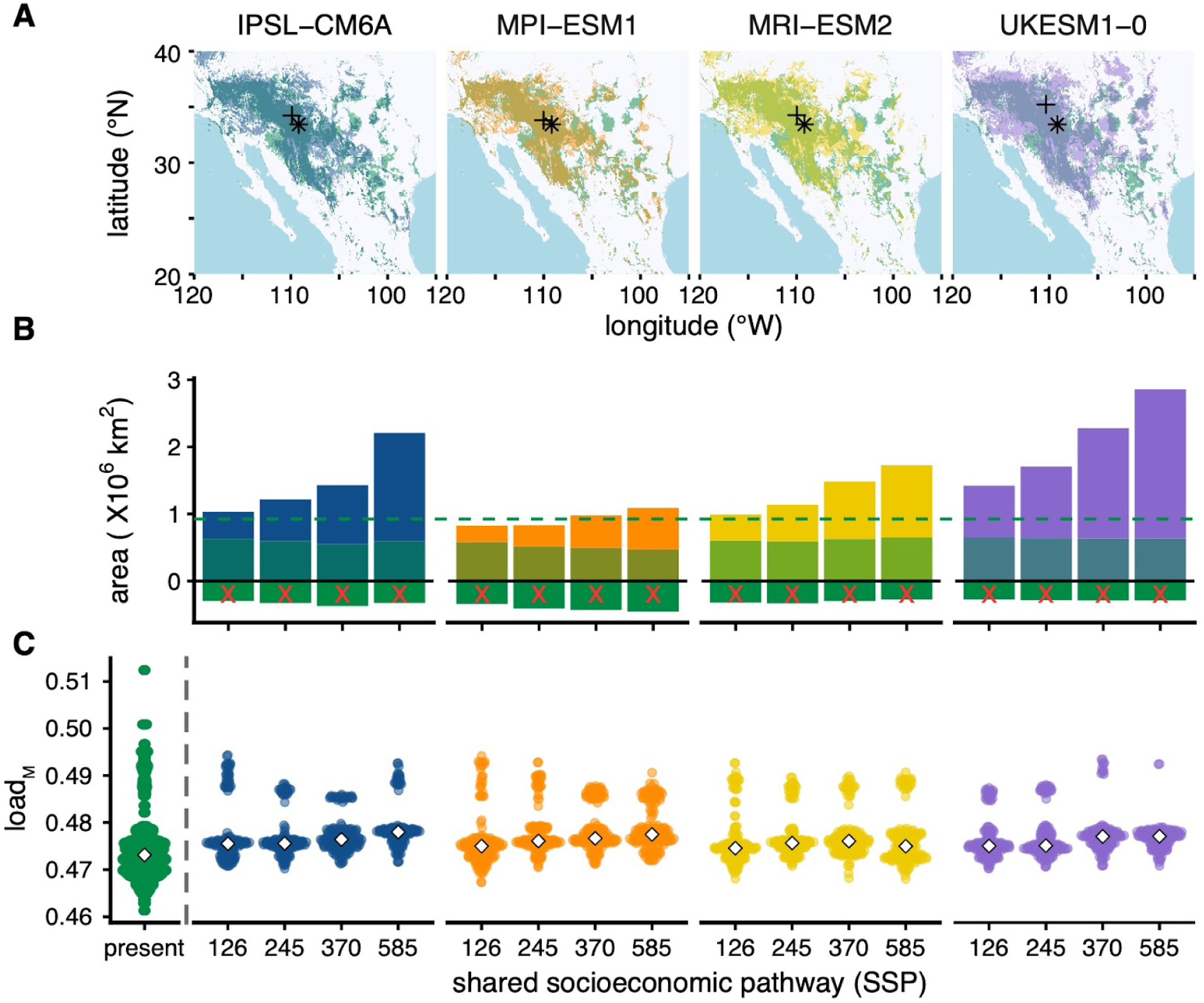
Predicted species distribution models and load_M_ in 2100. (A) Predicted species distribution models in 2100 by Earth Systems Model (ESM) for SSP126 (various colors) compared to the present SDM (green). Blue corresponds to IPSL-CM6A, orange is MPI-ESM1, yellow is MRI-ESM2, and purple is UKESM1-0. In each map, the predicted future geographic centroid is denoted by a plus (+), while the present geographic centroid is represented by an asterisk (*). Predicted SDMs for additional shared socioeconomic pathways are shown in **Figure S6**. (B) Predicted SDM area in 2100 by ESM and SSP. The height of each bar indicates the total predicted area subdivided by new areas that were not part of the present SDM (top), overlapping area between the predicted future and present SDM (middle), and the area of the present range predicted to be lost (green, negative values). The horizontal dotted green line indicates the area of the present SDM. (C) load_M_ predicted by the random forest model per each ESM and SSP combination. Diamonds represent median values.

Increasing emissions were associated with a greater degree of range shift, ranging from 160.91 to 348.57 km for SSP245, 238.01 to 510.44 km for SSP370, and 226.53 to 697.69 km for SSP585 (**Figure S6**). Potential species range area was predicted to increase under all scenarios except for the MPI-ESM1-2-HR ESM at SSP126 and SSP245. The change in range area was greater under increased emissions, with predicted changes of -10.8% to 53.8% by area for SSP126, by -10.2 to 84.5% for SSP245, by 5.87 to 146% for SSP370, and 17.8 to 209% for SSP585 (**Figure 4B**). The predicted range shifts included losses to the present range, particularly in the southernmost extreme.

By comparing the current and projected species distribution models, we estimated the minimum dispersal distance for each sample by 2100. As with our previous analysis, we only calculated dispersal for samples collected in regions occupying present but not future predicted range; we assumed that samples collected in overlapping regions,which represents the vast majority of samples, will not migrate (i.e. dispersal = 0). In total, we estimated that the lineages of between 1 (MRI-ESM2-0, SSP585) and 17 (UKESM1-0-LL, SSP585) individuals will need to disperse by 2100, with a minimum predicted migration between 2.41 and 306.55 km (median 6.19 Km).

Finally, we used the trained RF model to predict how load_M_ might change by 2100. As input to the model, we used the forecasted bioclimatic data along with species’ range features predicted by the future SDMs. In these analyses we kept H_O_ constant, because we had no basis to predict its values in the future. The RF models yielded two primary insights about load_M_ in 2100: first, load_M_ is generally predicted to increase for the majority of individuals and, second, variation in load_M_ is forecast to decrease markedly, with outliers regressing towards the mean (**Figure 4C**). Specifically, load_M_ is predicted to change by -5.16% to 3.33% per individual, with samples at more northern latitudes (latitude > 36) predicted to have reduced load_M_ while individuals closer to the present range centroid are expected to have increased load_M_.

## DISCUSSION

Understanding the relationship between climatic and genetic variation is essential for predicting which populations will thrive, persist, or face extinction in the future. Historically, the probability of persistence has been evaluated by estimating species distribution models (SDMs), but they have an important limitation: they ignore the ability for a species to evolve (Collart et al. 2020; Garzón et al. 2019). Newer methods have worked toward incorporating the potential for evolutionary change by using population genomic information in concert with climate predictions (Fitzpatrick & Keller 2015; Exposito-Alonso et al. 2019; Capblancq, Fitzpatrick, et al. 2020; Waldvogel et al. 2020). One such method relies first on identifying mutations that likely contribute to local adaptation and then on inferring the relationship between bioclimatic variables and the frequency of adaptive variants (Fitzpatrick & Keller 2015). The inferred relationship can then be used to predict future shifts in allele frequencies under projected climates, providing a measure of pre-adaptation to climate shifts. This and similar approaches have been applied to evaluate potential persistence of wild populations from several species [reviewed in (Capblancq, Fitzpatrick, et al. 2020)] and also to assess agronomic suitability of specific genotypes in projected climates (Rhoné et al. 2020; Aguirre-Liguori et al. 2022). These approaches do not, however, incorporate evolutionary processes beyond local adaptation, such as genetic drift, gene flow, migration and dispersal (Waldvogel et al. 2020). Another major limitation is that they usually [although not always (Bay et al. 2018; Ruegg et al. 2018; Booker et al. 2020; Rhoné et al. 2020)] focus on a tiny subset of genetic variants - i.e., identifiable, putatively adaptive variants - and thus ignore most genetic information.

An overarching challenge is how to include additional categories of genetic variation into climate models and also to evaluate whether the additional information has any predictive value for evaluating species’ persistence in the face of climate change. Here we have made a modest step toward this end by studying the relationship between a measure of deleterious variation (load_M_), species’ range dynamics, and climate in *V. arizonica*, a wild grape species endemic to the American Southwest. Our investigation of mutational load is relevant not only for evaluating potential persistence but also for a number of questions in evolutionary biology, such as potential differences between edge and non-edge populations (Vucetich & Waite 2003), relationships between genetic diversity and range centroids (Eckert et al. 2008; Lira-Noriega & Manthey 2014), and dissimilarities between leading versus trailing edges during range shifts.

Our study employed both population genomic and bioclimatic data. The former consisted of resequenced genomes from 162 individuals sampled throughout the predicted range (**Figure 1B**), and the latter was bioclimatic data from WorldClim that was used, in part, to estimate SDMs that became the basis for geographic distance measures. For example, we inferred a “present day” SDM (based on bioclimatic data averaged from 1970 to 2000) and estimated the distance of sampled locations from the center of the geographic range, the environmental niche and from the niche edge. We also estimated an SDM from the Last Glacial Maximum (LGM). A comparison between the two SDMs indicates that the current range of *V. arizonica* has expanded over the last ∼22,000 years, and it also implies that some of our sampling locations represent historical dispersal events, particularly in the North (**Figure 2A**). We also estimated SDMs based on climate projections to the year 2100, considering multiple climate models (**Figure 4A&B**, **Figure S6**). These future SDMs suggest that the range of *V. arizonica* will continue to expand, suggesting it is well situated to persist in the face of climate change, at least by this measure. *V. arizonica* is atypical in this respect, because a recent SDM-based study of 600 North American CWRs found that ∼85% are either vulnerable or endangered (Khoury et al. 2020).

### Measuring and interpreting mutational load

A comparison between the present-day and LGM SDMs imply that the Northern samples represent historical dispersal events at a leading edge of range expansion (**Figure 2A**). Given previous observations in plants that have found increased load in edge populations and/or reduced *Ne* in those populations (Willi et al. 2018; Takou et al. 2021; Willi et al. 2022; Sánchez-Castro et al. 2022; Cisternas-Fuentes & Koski 2023), it may not be too surprising that these samples had higher values of load_M_ (**Figure 1B)**. Given these load_M_ estimates, we asked two questions: i) Is load_M_ predictable across locations? ii) If so, what features are important for predicting load_M_?

We addressed the two questions using an RF framework, but here we focus on aspects and assumptions of our analyses before interpreting their outcomes. One drawback is that the reported feature importances may not reflect biological importance. Our impression is that this issue is often ignored in the biological literature; it frankly may not matter if one is only interested in predicted outcomes and not the features that contribute to that outcome. One potential solution to this problem is feature selection, which identifies a subset of features to use in modeling based on correlations, clustering analyses or other criteria. However, when the goal is to identify biologically relevant predictors, this approach has the potential to inadvertently remove important predictors - i.e., to toss out the veritable ‘baby with the bathwater’. Another approach is dimensionality reduction, which transforms correlated variables into orthogonal components. The challenge with dimensionality reduction is that it is difficult to interpret orthogonal components in terms of biological meaning. Here we used Johnson’s relative weights to retransform orthogonal vectors back onto the original features (Johnson 2000). Johnson’s relative-weight approach has been utilized widely across many scientific fields but rarely, as far as we could assess, in applications that rely on multifaceted biological and genomic data [e.g. (Core et al. 2014; Chen et al. 2018; Ghanipoor-Samami et al. 2018; Shen & Chen 2020; Li et al. 2022).]

Second, we relied on load_M_ as a measure of mutational load under the assumption it relates to genetic load. Measuring genetic load directly is notoriously difficult because it is a theoretical construct related to the unmeasurable (i.e., a decline in fitness relative to a theoretical fitness optimum) (Crow 1958). Here we used a measure, which we call load_M_, that summarizes the enrichment in the proportion of nSNPs (or dSNPs) to presumably neutral sSNPs (Renaut & Rieseberg 2015; Willi et al. 2018). Load_M_ is a desirable metric both because it measures the relative increase in nSNPs or dSNPs relative to a presumably neutral class (sSNPs) that captures aspects of population history, because it utilizes information from a large component of genomic variation and because it is analogous to well-used d_n_ / d_s_ analyses. It is not, however, without limitations. Like all measures of mutation load, it is strictly additive, assumes that each putatively deleterious allele has the same effect on fitness, and does not take into account potential dominance relationships between alleles that might obscure the fitness effects of deleterious mutations (i.e. masked load).

### Features that predict mutational load, including bioclimatic variables

We used RF to predict load_M_ based on 24 potentially informative features, including four geographic descriptors, one genetic descriptor, and 19 bioclimatic variables. The two RF implementations - i.e., with and without Johnson’s transformation - agreed on the high ranking of the relative importance of many predictors, including bioclimatic variables, dispersal distance, observed heterozygosity, and distance to the geographic centroid (**Figure 3B&D**). By contrast, the distances to the environmental niche centroid and to the geographic range edge had little predictive importance. We found the lack of importance explained by the niche centroid particularly surprising, because previous work has suggested that the distance from the niche centroid often better explains patterns of genetic diversity across a landscape than the distance to the geographic centroid. However, previous work has found that the relevance of the niche versus geographic centroids does vary among species and study systems (Lira-Noriega & Manthey 2014). We believe that distance to the niche centroid is a poor predictor in *V. arizonica*, because while it reflects the magnitude of difference from the niche centroid, it does not account for the direction of change. For instance, if the niche spans a cold-to-warm gradient, samples at either extreme could have identical values for niche distance despite being in vastly different environments. Thus, while distance from the niche centroid is not particularly predictive for *V. arizonica*, it may remain a useful feature for studying other species.

Our analyses lead to at least two general conclusions. First, although theory has established that mutation load is governed primarily by stochastic genetic drift, it is nonetheless also highly predictable based on the combination of features in our analyses; the RF models yielded R^2^ > 0.60 on withheld test data (**Figure 3A&C**). Second, several bioclimatic variables were among the top-ranked predictors. The important climate predictors were biased toward precipitation (e.g., bio8, bio12 to bio19) compared to temperature-specific variables (bio1 to bio7 and bio9 to bio11) (**Figure 3B&D**). There was some disagreement in the ranks of relative importances for the two RF implementations; the most obvious differences were the ranked importances of bio8 and dispersal, although both were among the top-ranked variables in both implementations. A concern is that features might be predictive because they covary with genetic relatedness. But genetic relatedness does not fully explain the importance of bio8, for example, because it is a highly significant (p = 2.40 X 10^-13^, **Table S1**) predictor in univariate mixed linear models that include genetic relatedness as a cofactor.

To be clear, we are not arguing that bio8 or any of the other important bioclimatic conditions affect genetic load directly. Instead we suspect that they reflect ecological conditions that affect population size, population density or perhaps additional aspects of population history that are not fully captured by genetic and geographic predictors (Willi et al. 2018). Genetic predictors are known to be important based on much previous work [review in (Robinson et al. 2023)], a fact reasserted but by H_o_ and dispersal distance being strong predictors of load_M_ in *V. arizonica* (**Figure 3B&D**). More generally, we do not yet know if climatic variables will be similarly useful for predicting mutational load in other systems. However, we suspect that they will be, based on well known relationships among range limits, marginal habitats, life history traits, and genetic diversity.

### Load_M_, fitness, and projections into the future

The ability to predict load_M_ in the present is important for two key reasons. First, as previously noted, it offers insights into the relative importance of factors contributing to the accumulation of genetic load. Second, it provides a foundation for projecting load_M_ into the future. We recognize that such projections may seem ill-conceived, given that mutational load is largely shaped by random genetic drift and hence likely to be inherently difficult (or even impossible) to predict with precision. In contrast to load, adaptive variants can be modeled as a component of the deterministic process of selection (Fitzpatrick & Keller 2015; Capblancq, Morin, et al. 2020; Gain et al. 2023). However, here we have shown that load can be modeled accurately in the present, through features that provide a credible basis for future projections (Aguirre-Liguori et al. 2021). This approach paves the way to incorporate deleterious variants (and not just adaptive variants) to evaluate the fate of species in the face of climate change.

As an illustration, we utilized data from projected climates (to the year 2100). We used each SDM to update features (e.g., bioclimatic variables and distance features) for the RF model. Using 16 discrete climate models based on four ESMs and four SSPs, we forecasted consistently higher mean load_M_ at the end of this century **(Figure 4C)**. Each of the 16 projections also had reduced variance of load_M_ [Var(load_M_)] across samples. We suspect the lower variance has three causes. First, the predictive models are unlikely to fully incorporate evolutionary processes, like genetic drift and gene flow, that may contribute substantially to Var(load_M_). Thus, Var(load_M_) may be systematically under-predicted by our methods. One potential path forward to this problem is generating training data from spatially explicit population genetic simulations that include all relevant evolutionary processes. At present, however, spatially explicit simulations require simplifications and assumptions that are likely to make simulated data an inaccurate representation of reality (Battey et al. 2020; Láruson et al. 2022). A second cause of the reduced (load_M_) could reflect limits in the use of RF for extrapolation. A final explanation for the reduction in Var(load_M_) relates to predicted shifts in the range of *V. arizonica* over time. As this shift occurs, our sampled edge locations are expected to no longer be at a leading edge where load_M_ is likely to be most extreme.

Our focus on mutational load is only meaningful if load_M_ reflects a component of fitness; a relationship between mutational load and fitness is commonly assumed but rarely tested [but see (Mezmouk & Ross-Ibarra 2014; Sánchez-Castro et al. 2022)]. Unfortunately, we cannot test this relationship directly in *V. arizonica* because we do not have fitness measures of our genotypes in the wild. However, the same genotypes have been phenotyped in the greenhouse for two agronomic traits: chloride exclusion, a measure of salt tolerance (Heinitz et al. 2020), and a quantitative measure of resistance to *X. fastidiosa* (Riaz et al. 2020; Morales-Cruz et al. 2023), the causative agent of PD. We contrasted these phenotypes to load_M_ and found no evidence that mutational load is related to chloride exclusion phenotypes. However, load_M_ is correlated with assayed concentrations of *X. fastidiosa* post-infection (R^2^ = 0.33; p = 1.5 X 10^-4^, **Figure 5**). In other words, plants with higher load_M_ are more susceptible to PD.

**Figure 5.**
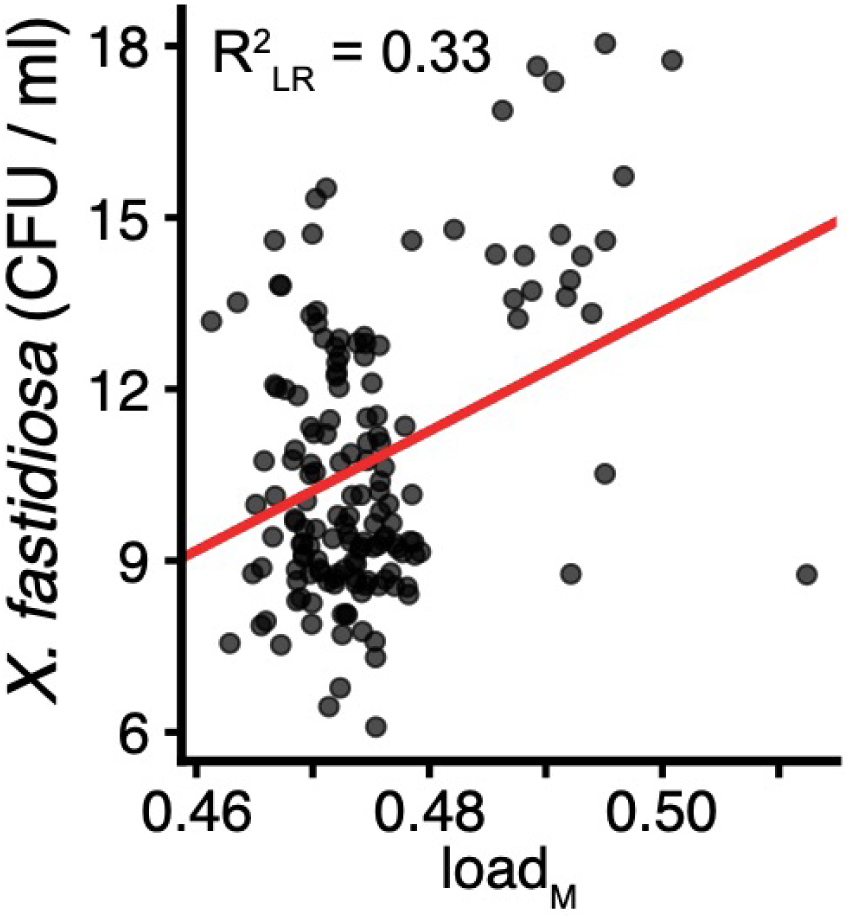
Relationship between load_M_ and *X. fastidiosa* resistance. Scatter plot of *X. fastidiosa* (XF) concentration in a greenhouse experiment against load_M_. The red line indicates fixed effects from the linear mixed model.

One reasonable interpretation is that plants with higher load_M_ have lower fitness and thus are more susceptible to disease. We suspect, however, that this interpretation is not complete, for two reasons. First, *V. arizonica* resistance to PD has been mapped to a few key genomic regions containing disease resistance genes (Morales-Cruz et al. 2023). It thus seems unlikely that a genome-wide measure of genetic variability (i.e., load_M_) directly reflects a major component of disease susceptibility. Instead, we point out that bio8 is an important feature for predicting both load_M_ (**Figure 3D**) and PD susceptibility. In *V. arizonica* resistance to the bacterium is less common below a threshold of bio8 < 10 °C; infections are rarely observed below this threshold in crop species (Morales-Cruz et al. 2023). We previously proposed a selective interpretation for higher disease susceptibility of *V. arizonica* at bio8 < 10 °C, which is that: i) *X. fastidiosa* is not present where bio8 < 10 °C, ii) resistance incurs a cost in the absence of pathogen pressure, and iii) as a result, resistance is selected against in environments where bio8 <10 °C (Morales-Cruz et al. 2023). Our analyses here suggest an additional evolutionary interpretation: the loss of disease resistance may be a manifestation of the accumulation of deleterious variants in Northern edge populations.

### Lessons for *V. arizonica* and beyond

This study was designed to address general questions about mutational load, to explore whether load correlates with climatic data, and to move toward using this information for interpreting potential persistence in climate change. While our questions were general, we focused on *V. arizonica* as a convenient model and as an example of a valuable CWR. Overall, *V. arizonica* seems generally well-situated to adapt to climate change based on our analyses. SDMs predict continued expansion of its range across hot and dry environments (**Figure 4A**, **Figure S6**) and predicted increases of genetic load are only moderate (**Figure 4C**).

Even if these observations are accurate, they do not necessarily imply that *V. arizonica* is poised to thrive in the future. One major question is whether the species will be able to disperse effectively to take advantage of an expanding niche. *V. arizonica* is consumed by birds and mammals that can, in theory, disperse seed to suitable habitats as the niche expands over time. However, dispersion is difficult to predict across heterogeneous landscapes, and this factor needs to be included more often in predictive models (Razgour et al. 2019; Aguirre-Liguori et al. 2021). *V. arizonica* has another potential evolutionary advantage, which is that it readily hybridizes with other *Vitis* species. Hybridization can fuel rapid adaptation in *Vitis* and other plant genera (Pease et al. 2016; Morales-Cruz et al. 2021), but this addresses another important point: species do not evolve in isolation. Range is often limited by biotic interactions including closely-related species, pollinators, dispersers, pathogens and pathogen vectors. These biotic interactions are rarely captured in climate projections of species’ persistence. There is thus an urgent need to assess range and climate simultaneously across multiple interacting species, just as there is a need to consider non-adaptive genetic variation.

Additional uncertainties apply across plant species. Range expansion often favors a shift to a selfing mating system in edge populations, likely bolstering reproductive success at low population densities (Koski et al. 2019). This implies that species with flexible mating systems may be better equipped to expand their range. However, *Vitis* species possess a highly-conserved, dioecious sex determination system that ensures outcrossing. Interestingly, this mating system can produce hermaphrodites via a simple recombination event between male and female alleles, as occurred during grapevine domestication (Massonnet et al. 2020), but hermaphrodites are rarely observed in nature, suggesting that they are strongly selected against. This raises the question: Will the evolutionary conservation of the dioecious mating system limit the potential for range expansion for *V. arizonica*? Additionally, while our analyses have largely focused on individuals at the leading edge of expansion, we have given less attention to the populations at the lagging edges, where suitable niches are disappearing. In these populations, dispersal, along with the rate and possibility of evolutionary rescue, are both crucial for persistence; the likelihood of rescue depends on population attributes and the severity of habitat deterioration (Bell 2017) and possibly the extent of genetic load.

## MATERIALS AND METHODS

### Variant calling

Illumina sequencing reads from NCBI BioProjects PRJNA731597 (Morales-Cruz et al. 2021) and PRJNA842753 (Morales-Cruz et al. 2023) were processed using trimmomatic 0.39 (Bolger et al. 2014) to remove adapters and low quality sequence using the following parameter string: “ILLUMINACLIP:"$ADAPTERSPE":2:30:10 LEADING:3 TRAILING:3 SLIDINGWINDOW:4:20 MINLEN:60”. Processed reads were then mapped to the *Vitis arizonica* B40-14 v. 2.0 genome assembly (Morales-Cruz et al. 2023) with bwa-mem 0.7.12r1039 (Li 2013) using the default parameters. SAM alignments were sorted and converted to indexed BAM using samtools 1.17 (Danecek et al. 2021). Sequencing duplicates were marked in the alignments using the picard MarkDuplicates module included in GATK 4.2.6.1 (Van der Auwera & O’Connor 2020).

SNPs were called using GATK HaplotypeCaller in GVCF mode followed by joint calling across all samples with GenotypeGVCFs. Bcftools 1.17 (Danecek et al. 2021) was used to filter SNPs, keeping only biallelic SNPs with quality of 20 or greater that also passed the GATK “best practices” hard filters: excluding sites with “QD < 2 | FS > 60 | SOR > 3 | MQ < 40 | MQRankSum < -12.5 | ReadPosRankSum < -8.0.” Sites with minor allele frequency less than 0.01, with site depth greater than the mean plus standard deviation of depth across all sites, and that had more than 5% missing calls between individuals were filtered. Annotation for putative SNP effects was done with SIFT-4G (Vaser et al. 2016) using a custom database based on the *Vitis arizonica* B40-14 v. 2.0 genome assembly and annotation (Morales-Cruz et al. 2023). The SNPs were polarized by including 6 outgroup samples consisting of 3 individuals each from *Vitis girdiana* (SC11, SC33, and SC51) and *Vitis monticola* (C20-93A, T_03-02_S01, and T40). Only sites with consistent homozygous genotypes and no missing data across outgroup samples were used for subsequent analyses.

As a final filtering step, principal component analysis (PCA) on SNPs was done using only the *V. arizonica* samples using plink 2.0 (Chang et al. 2015). Variants were pruned for linkage disequilibrium in 50 variant windows, with a step size of 10, and R^2^ threshold of 0.20 prior to the PCA. The 10 samples on the extremes of PC 1 in the SNP PCA per the 1.5 IQR rule (**Figure S1**) were removed from all subsequent analyses.

### Estimation of mutation load

Mutational load (load_M_) was estimated from the polarized SNP calls for each individual similarly to (Willi et al. 2018), but for single individuals and considering counts of alleles instead of homozygous sites. Briefly, mutational load was calculated as 𝑃𝑛 / (𝑃𝑛 + 𝑃𝑠), where P_n_ corresponds to the proportion of derived nSNP calls at nonsynonymous sites and P_s_ corresponds to the proportion of derived sSNPs calls at synonymous sites. A complementary measure of mutational load based on the subset of nonsynonymous SNPs annotated as putatively deleterious was also calculated: 𝑃𝑑 / (𝑃𝑛 + 𝑃𝑠), where P_d_ was the proportion of derived deleterious SNPs calls at deleterious sites. Although we employed load_M_ measures with precedent, we also investigated the behavior of alternative measures. We considered, for example, the simpler measure 𝑃𝑛 / 𝑃𝑠 which is essentially π^N^/ π^S^, but this simpler measure was essentially identical (R^2^ > 0.99) to the Willi et al. (2008) measure.

### Species distribution modeling

Species distribution models for *V. arizonica* were calculated for both the present and the Last Glacial Maximum following the procedure described in (Aguirre-Liguori et al. 2022). To assemble the data for constructing the SDMs, the WorldClim 2 bioclimatic variables (2.5 minute resolution, mean of observations from 1970 to 2000) were extracted for all available geographic references of *V. arizonica* from the Global Biodiversity Information Facility ((Occdownload Gbif.Org 2020)) (accessed 2022-06-06) and for the sampling locations (Table S1) using the raster 3.6-26 R package (Hijmans 2023). The CoordinateCleaner 3.0.1 R package (Zizka et al. 2019) was used to remove duplicate references, as well as references that were outliers (locations that are at the top 5% quantile distribution of the mean distance to all other locations), present in water (and thus obviously incorrect), present at GBIF headquarters facilities, and locations that are within the vicinity of the centroid coordinates of countries or their capitals.

To build the SDM, first the bioclim correlated variables were pruned based on variance inflation factor, retaining variables with R < 0.8. Next, the background area was set by selecting the overlap between the pruned occurrence records and the terrestrial ecoregions defined by (Olson et al. 2001). The model was then built using the BIOMOD2 4.2-4 R package (Thuiller et al. 2009) and the Maxent algorithm (Phillips et al. 2006; Phillips & Dudík 2008) using 20 bootstrap replicates, utilizing 70% of occurrences as the training data and retaining 30% as a test dataset. The final distribution model was selected by evaluating True Skill Statistics among 10-fold internal cross-validation. To estimate how the distribution of *V. arizonica* has changed from the past to the present, and project how it will change from the present to the future, we projected the SDM using the same set of bioclimatic variables for the LGM layer (∼22,000 ya) and to the future bioclimatic layers described below. For all SDM models, all range areas were calculated using the expanse function from the terra 1.7-55 R package (Hijmans 2024), and the distance between centroids was calculated using the distHaversine function from the geosphere 1.5-18 R package (Hijmans 2022).

### Features

The features for prediction consisted of 24 variables summarized in Table 1. The 19 WorldClim 2 bioclimatic variables (2.5 minute resolution) were extracted for each individual by collection site coordinate using the raster 3.6-26 R package (Hijmans 2023). Observed heterozygosity (H_O_) was calculated from the SNP dataset after pruning for linkage disequilibrium (50 SNP windows, 10 SNP step size, R^2^ threshold 0.20) using plink 2.0 (Chang et al. 2015). The SDMs were used to generate the remaining predictors as described below.

The geographic centroid was defined for the SDM as the median coordinate of the present SDM. For each of the samples, the distance to the geographic centroid was then calculated as the Euclidean distance between each sampling location and the centroid. The distance to geographic range edge was calculated by using the distGeo function from the geosphere 1.5-18 R package (Hijmans 2022) to determine the distance between each sampling location and the nearest edge of the species range defined by the boundaries function implemented in the terra 1.7-55 R package (Hijmans 2024).

The distance to the niche centroid for each individual was calculated following the approximation of (Lira-Noriega & Manthey 2014). Briefly, the 19 WorldClim 2 bioclimatic variables (2.5 minute resolution) (Fick & Hijmans 2017) were extracted for all pixels in the present SDM. Next, a PCA of the bioclimatic data was performed and the first 6 PCs (explaining 95% of the variation in the dataset) were retained. The niche centroid was defined as the mean value among all observations along the 6 PCs. Finally, the distance to the niche centroid was then calculated as the Euclidean distance in multi-dimensional PC space between the niche centroid and the observation for each individual.

The final predictor was the estimated dispersal distance from the LGM to the present. The assumption of this calculation was that individuals with collection sites in present but not past range must have dispersed to their present locations since the last epoch while individuals present in the overlapping range did not necessarily need to disperse to their present location (dispersal = 0). Therefore, the geographic dispersal distance for each individual was calculated using the distGeo function from geosphere 1.5-18 R package (Hijmans 2022) between the collection site and the closest boundary of the past distribution.

### Statistical modeling

Random forest regression (RF) models (Breiman 2001) were built using the tidymodels 1.1.1 framework (Kuhn & Wickham 2020) utilizing the ranger 0.16.0 engine (Wright & Ziegler 2017) running in the R programming environment (R Core Team 2023). The dataset was split, allocating 75% of sample observations to training and reserving 25% for testing.

Hyperparameter optimization for mtry (the number of randomly sampled predictors used to split the decision trees), min_n (the number of observations required for a tree node to be split again after segmentation), and trees (the number of decision trees to be included in the ensemble) was conducted over the respective ranges of (1,20), (1,10), and (500,1000) through Latin hypercube sampling, selecting 100 unique combinations for evaluation. Optimal hyperparameters were determined via 10-fold cross-validation on the training data and were used for the final model fits (**Figure S4** and **Figure S5**). The permutation approach was used to calculate predictor importance. A static seed was used for all modeling to ensure reproducibility.

The data transformation to reduce collinearity between independent variables was done broadly following (Johnson 2000), wherein an orthogonal approximation (***Z***) of the original data (***X***) was generated and used to fit the model, with the resulting model coefficients (i.e. variable importance) then transformed back into the original data space for interpretation. Specifically, the training data was first scaled before singular value decomposition (SVD) was performed on the dataset. The data was then orthogonally transformed by the following formula: ***Z*** = ***P Q^T^***, where ***P*** represents the left singular vectors and ***Q*** represents the right singular vectors. The transformation parameter, **λ**, was then calculated by the following formula: **λ** = ***Q D Q^T^***, where ***D*** is a diagonal matrix with the singular values. In the context of RF models, the importance values from the model trained with the orthogonalized data were then approximated back into the original data space by multiplying column-wide normalized **λ^2^** by the importance values. To use the models trained on orthogonally approximated data for predictions, test data (***X_1_***) was first scaled using the column means and standard deviations of the original training dataset before being projected into the SVD space using the following formula: ***Z_test_***= ***X_1 scaled_ Q D^-1^* Q^T^** .

We also examined the data using linear mixed-models, which were built using the lmekin function from the coxme 2.2-18.1 R package (Therneau 2022). All predictors were scaled using the base::scale function in R (R Core Team 2023) prior to model fitting. A standardized relatedness matrix was calculated with gemma 0.98.5 (Zhou & Stephens 2012) and included as a random effect in the mixed linear models.

Multiple linear regression models were built using the tidymodels 1.1.1 framework (Kuhn & Wickham 2020) utilizing the lm engine included in the base stats R package (R Core Team 2023).

### Mantel test for correlation between genetic and climatic distance

Genetic distance (Hamming) based on all SNPs was calculated using plink 1.9 (Chang et al. 2015). Euclidean distance was calculated between climate variables using the dist function after standardization using the scale function in base R (R Core Team 2023). Mantel test was calculated using mantel.rtest function from the ade4 1.7-22 R package (Dray & Dufour 2007) using 1,000 replicates.

### Projections of climate and mutational load in 2100

We used the SDMs described previously and projected the future distribution models using the bioclimatic data for four Earth System Models (GFDL-ESM4, IPSL-CM6A-LR, MPI-ESM1-2-HR, MRI-ESM2-0, UKESM1-0-LL) representing four shared socioeconomic pathways (SSP126, SSP245, SSP370, and SSP585) for the period 2081-2100 downloaded from CMIP6 project (Eyring et al. 2016). The distance to geographic centroids and distance to range edge for each model were calculated as described previously. To calculate future dispersal, we followed the same approximation as above. First we evaluated if the location was covered by the present range and is predicted to be part of the future forecasted range. If so, we set the future dispersal distance to zero. If not, we estimated the geographic distance between the current location and the closest boundary of the nearest forecasted range edge. For all the locations in the future (locations that did not disperse, or the locations at the nearest future edge), we obtained the bioclimatic variables in the 16 future models described above.

The random forest models built using the data projected into orthogonalized space were used for all projections of genomic load into the future. As independent variables, WorldClim 2.1 bioclimatic variable projections for 2081-2100 (2.5 minute resolution) (Fick & Hijmans 2017) for all 16 ESM:SSP combinations were used, along with the projections of future geographic centroids, distance to geographic range edges, and distance to niche centroid. For dispersal, the values calculated previously (i.e. dispersal from LGM to present) were added to the projected dispersal into the future. Observed heterozygosity was kept constant.

### Phenotypic analyses

Mixed-linear models to detect association between phenotype and load_M_ were built using the lmekin function from the coxme 2.2-18.1 R package (Therneau 2022) using a standardized relatedness matrix calculated with gemma 0.98.5 (Zhou & Stephens 2012) as a random effect as described previously. The likelihood-ratio pseudo R^2^ was estimated using the r.squaredLR function from the MuMIn 1.48.4 R package (Bartoń 2024).

## DATA AVAILABILITY

All sequencing data referenced in this work is available from NCBI: BioProjects PRJNA731597 (Morales-Cruz et al. 2021) and PRJNA842753 (Morales-Cruz et al. 2023). The phenotyping data was also previously published (Heinitz et al. 2020; Morales-Cruz et al. 2023). All other data needed to reproduce the analyses presented here is present within the manuscript and supplementary material.

## ACKNOWLEDGEMENTS

We thank Edwin Solares for supporting the Gaut lab’s high-performance computing infrastructure and Rebecca Gaut for assisting with the initial generation of the sequencing data. BG is supported by NSF grant IOS-2323123.

