## Supplemental Figures for "Climate, population size, and dispersal influences mutational load across the landscape in *Vitis arizonica*"

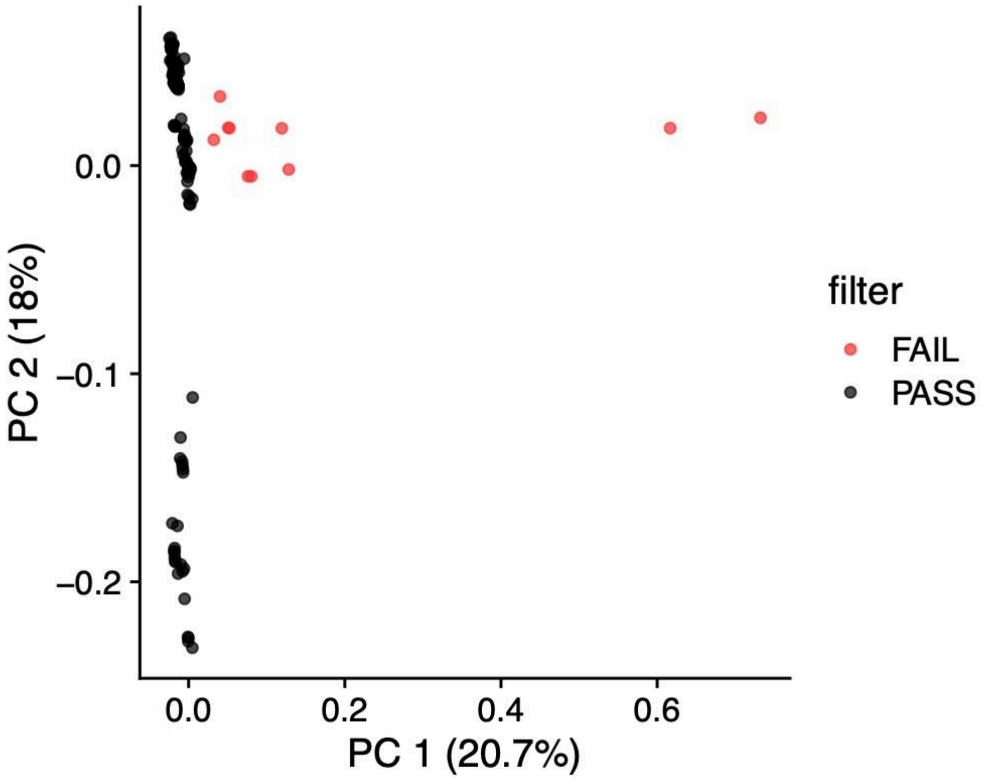


**Figure S1. Principal component analysis of SNP dataset**

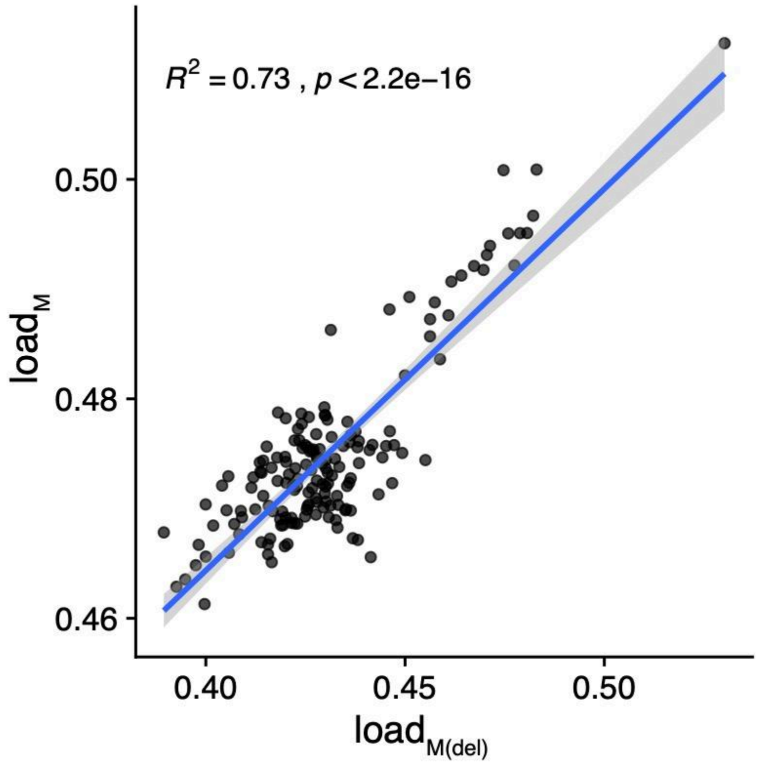


**Figure S2. Relationship between $load_M$ calculated with nonsynonymous compared to deleterious SNPs**

Blue line indicates linear model fit and gray shaded region is 95% confidence interval.

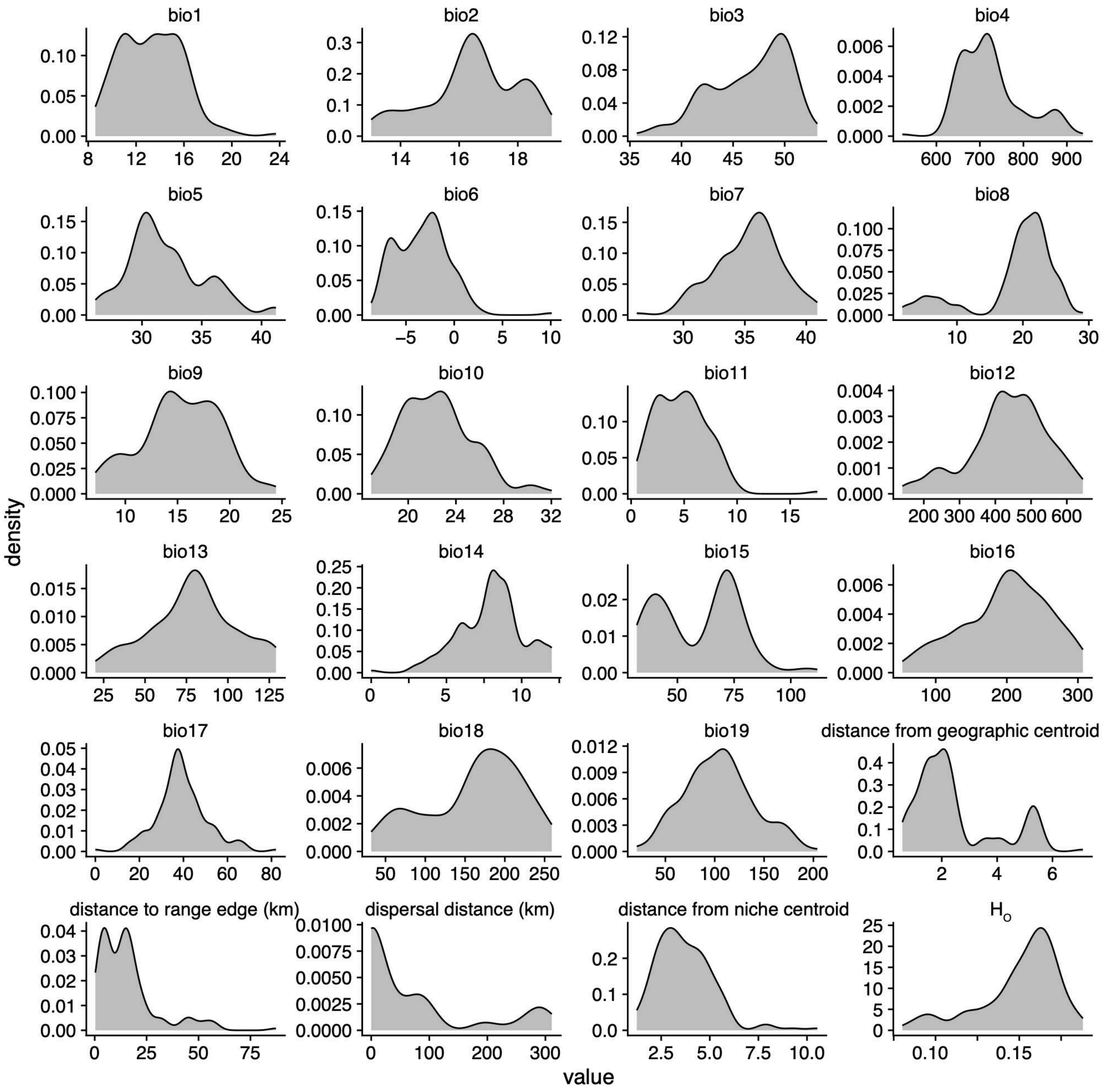


**Figure S3. Distributions of all features**

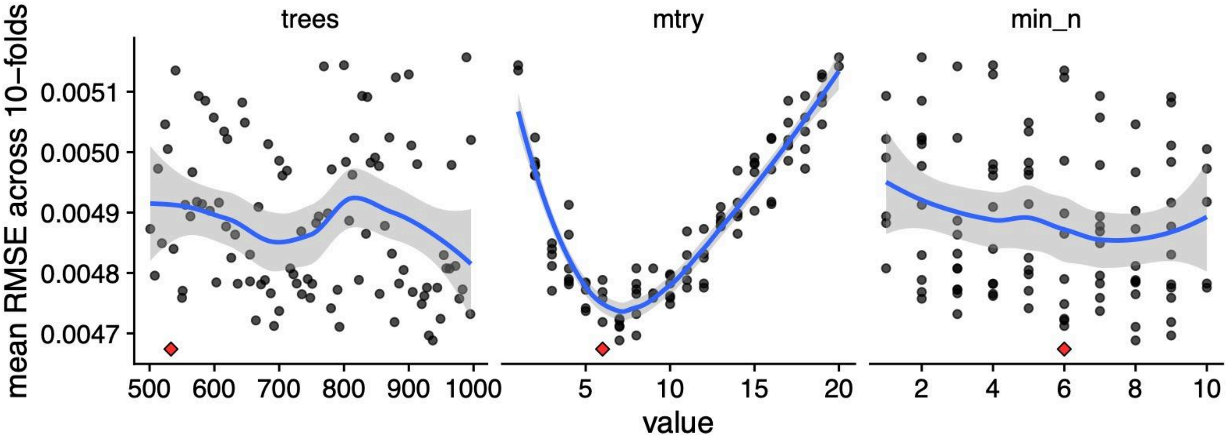


**Figure S4. Hyperparameter tuning for random forest regression model to predict load$_M$**
Blue line indicates smoothed conditional means using loess function while gray shading is 95% confidence interval. Red diamond denotes hyperparameter combination with lowest RMSE.

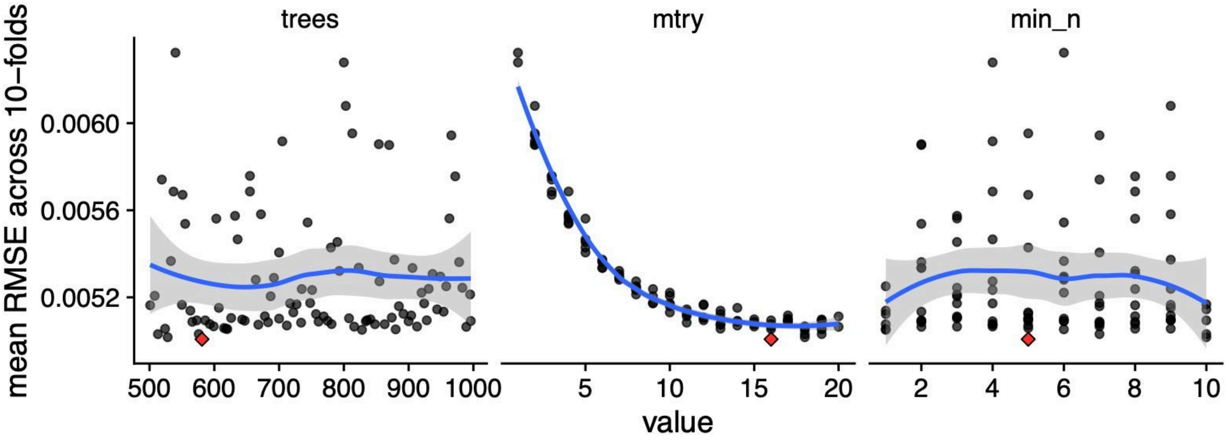


**Figure S5. Hyperparameter tuning for random forest regression model to predict load$_M$ (transformed data)**

Blue line indicates smoothed conditional means using loess function while gray shading is 95% confidence interval. Red diamond denotes hyperparameter combination with lowest RMSE.

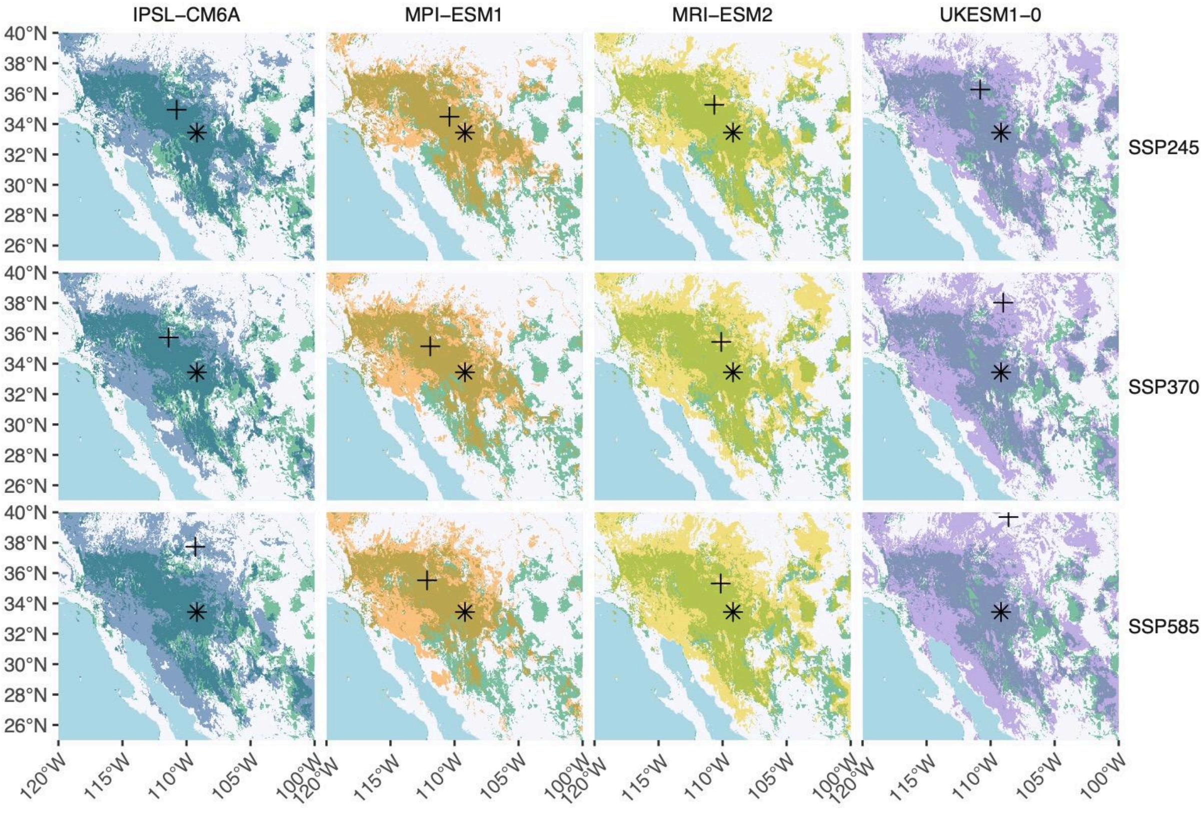


**Figure S6. Current vs. predicted species distribution models in 2100.**
Plus is predicted future centroid and star is current centroid. Green is the present species distribution and other colors are the future models.
